# Cortico-striatal dynamics across working memory stages

**DOI:** 10.1101/2025.07.04.663202

**Authors:** Maxime Villet, Benjamin Azoulay, Jacques Barik, Hélène Marie, Ingrid Bethus

## Abstract

Working memory depends on the temporary retention and manipulation of information, bridging the gap between short-term memory and information processing functions. However, when the same working memory task is repeated over several days, it raises the question of whether the rule or task set becomes automated (or proceduralized). The medial prefrontal cortex (mPFC) is crucial for working memory. Yet, the role of the dorsolateral striatum (DLS) in the automation (proceduralization) of rules or task sets remains to be clarified. Using a longitudinal approach of the “delay non-match to place” (DNMP) task in a T-maze combined to chemogenetic inhibition of the mPFC or DLS in mice, we show that the mPFC becomes less critical in the maintenance phase of the task as behaviour progressively shifts toward automation. During this phase, the DLS facilitates automated processing. Accordingly, silencing through chemogenetic inhibition of the DLS during maintenance triggers an adaptation in learning strategies, reactivating a goal-directed behaviour. Our findings strengthen memory traces as a dynamic reorganization of neural networks, challenging the classical view of information migration between brain structures. We here propose that the memory trace remains in a dormant state—less energy-consuming for the system—while still allowing for rapid flexibility in case of task modification.

## INTRODUCTION

Working memory refers to the process of temporarily holding new incoming information, while simultaneously performing cognitive manipulation on this information for a short duration (Eichenbaum, 2002). Alan Baddeley and his colleagues first recognized the importance of distinguishing the cognitive and storage processes in short term memory, and replaced the concept of a unitary short-term memory with a multiple components conception of working memory that mediates both storage and central executive functions (Baddeley, 2000, 1986; Baddeley and Hitch, 1974).

However, in this multiple component’s conception of the working memory, less attention has been put on the question of long-term storage. Indeed, working memory cannot be reduced to a short-term memory process. During learning of a working memory task, certain aspects, labelled task-set such as the rules, expected stimuli, and motor responses can become routine and may be consolidated into a long-term memory such as an automated procedural memory (Monsell and Graham, 2021; Sakai, 2008). Let’s take the example of a chess player. Each game is different, and the sequence of actions to checkmate the opponent will be different, but the rules of the game, the expected stimuli and the motor responses remain the same from game to game. In our context, although the complexity of the cognitive task assigned to the mice is not comparable to that of chess, we still anticipate the development of a rule. In some studies, this procedural memory is referred to as “reference memory” in animals, which stores the rule needed to succeed during the task (Miller, 2013; Yoon et al., 2008). This aspect may explain why, through daily training in a working memory task, we observe an improvement of the performance associated with a reduction of the time response, reflecting a more efficient processing of the rule contained in procedural memory (Logie et al., 2020; Oberauer, 2006; Vandierendonck, 2016).

When analyzing the structures that support working memory, most studies aim to untangle the three components of short-term memory dynamics within a single trial: encoding, maintenance, and retrieval. Spellman et al. (Spellman et al., 2015) found that direct inputs from the ventral hippocampus (vHPC) to the medial prefrontal cortex (mPFC) are essential for successfully encoding task-related cues in a spatial working memory task. In contrast, Wilhelm et al. (Wilhelm et al., 2023) identified the direct projection from the mPFC to the dorsomedial striatum as crucial for maintaining spatial working memory and Akhlaghpour et al. (Akhlaghpour et al., 2016) found that neurons from the dorsomedial striatum were sequentially activated throughout the course of the delay period.

Broadly, the involvement of mPFC has been described during each of these phases (Vogel et al., 2022), classifying it as a key structure supporting the short-term component (Uylings et al., 2003; Yang et al., 2014) and executive functions such as decision-making, attention, and behavioral flexibility of working memory (Delatour and Gisquet-Verrier, 2000; Euston et al., 2012; Gisquet-Verrier and Delatour, 2006; Ragozzino et al., 1999). Regarding the mPFC subregions, Killcross et al. (Killcross, 2003) distinguish between the prelimbic mPFC, which is responsible for the performance of voluntary response (goal-directed), and the infralimbic mPFC, which is involved in the incremental ability to develop habits that are no longer voluntary or goal-directed via prolonged training.

However, few studies address the question of the long-term storage of working memory after overtraining. Yoon et al. (Yoon et al., 2008) demonstrated the role of the dorsal hippocampus in the “reference memory” versus the mPFC in choice accuracy in a working memory task (Euston et al., 2012).

Although the role of the dorsolateral striatum (DLS), is well-documented for its role in inflexible automated behaviour (Balleine and O’Doherty, 2010; Hilario et al., 2012a; Packard and Knowlton, 2002; Yin et al., 2004), its implication in the context of “automated procedural memory” or “reference memory” in working memory tasks remains elusive. Thus, here, we tested whether the DLS could be at play in this context. To test this hypothesis, we trained mice in a delayed non-match-to-place (DNMP) spatial working memory task in a T-maze and discriminated between two distinct phases of memory, which are the learning and the maintenance phases. Then, using chemogenetic tools, we silenced neurons of the mPFC or DLS during either the learning phase or the maintenance phase and examined its outcomes on memory. Our results showed that, as expected, the mPFC is key for learning, but surprisingly its inhibition during maintenance had no effect on memory performance. In contrast, inhibition of the DLS affected maintenance but not learning. Using a devaluation test, we demonstrated that the learning phase is goal-directed, while the maintenance phase operates automatically and is independent of reward value. Interestingly, inhibiting the DLS during the maintenance phase did not affect memory performance potentially because of the re-engagement of goal-directed behaviour under the control of the mPFC. This possibility was further supported by the observation that simultaneous inhibition of the mPFC and DLS during the maintenance phase impaired memory performance. Thus, we show that there is a reorganization of brain circuits, with the possibility of dormant memory traces in latent circuits that can be instantly reactivated as soon as the secondary circuit, which has taken over, fails.

## METHODS

### Animals

Experiments were performed on forty-eight C57BL/6JRj male mice (Janvier Labs, France) aged 7 weeks at the beginning of experiments. Animals were housed in groups of 6 in a standard laboratory animal house. The experiments were carried out with precise control of the diurnal cycle (12h L:D without inversion) and temperature (fixed at 22 °C). Animals had free access to food and water until the beginning of protocol. All procedures were carried out in accordance with the recommendations of the European Commission (2010/63/EU) for care and use of laboratory animals and approved by the French National Ethical Committee (APAFIS#29469; E061525).

### Stereotaxic virus injection and clozapine-n-oxide treatment

We transduced an AAV virus expressing the Designer Receptor Exclusively Activated by Designer Drugs (DREADD) hM4Di receptor that inhibits neuronal firing when activated by the designer drug CNO, together with mCherry. A modified clozapine N-oxide (CNO)-activated human muscarinic receptor, hM4Di (Armbruster et al., 2007), was selectively expressed with a red fluorescent protein (mCherry) in either the mPFC or the DLS, or both, using an adeno-associated viral expression system (AAV8-hM4Di-hSyn-mcherry) from the University of North Carolina vector core facility, USA. Three weeks before behavioural experiments, the virus was injected in the brain by stereotaxic injection. This surgery was performed using a stereotaxic frame (Kopf Instruments) under general anesthesia with intraperitoneal (i.p.) xylazine and ketamine (10 mg/kg and 150 mg/kg, respectively; Centravet, France). The virus (titration 10E12 particles/ml) was injected bilaterally at 150 nl per site in the mPFC (AP: +1,7 mm; ML: + 0,27 mm; DV: -2,5 mm /-2 mm from bregma) and/or 350 nl per site in the DLS (AP: +0.6 mm /+1 mm; ML: + 2.25 mm; DV: -3 mm) with a flow rate of 100 nl/min. Cannulas were left in place for another 5 min to avoid backflow.

CNO (Enzo Life, France) was prepared in saline solution 0.9% NaCl at 1mg/kg and injected i.p. 30 min before the behavioural tasks to activate hM4Di and thus silence virally-transduced neurons. Control animals were injected with Saline (10ml/kg).

### Histology

At the end of behavioural protocols, mice were sacrificed by intra-aortic perfusion of 4% paraformaldehyde under deep anesthesia for brain collection and localization of mCherry-tagged cells. Brains were then cut coronally with a vibratome (Microm) at 40 μm. Slices were stained 10 minutes with DAPI (1:10000, #D1306, Invitrogen) and mounted in Vectashield solution (H-1000, Vector Laboratories). Slices were imaged with the Vectra 3 slide scanner (Akoya Bioscences, USA) using a 10x/0.2 objective. Animals not showing adequate injection (localization or expression) were removed from the study (14 mice).

### Delayed no-match to place (DNMP)

All behavioural tasks were performed in a T-maze apparatus. This maze consisted of a starting box containing clean bedding and closed by a removable door. It had a central stem of 35 cm long and two arms perpendicular at the end of the central stem, each of 35 cm length. The width of the arms was 5 cm. The walls of the T-maze were made of transparent Plexiglas allowing the animal to see external cues. At each end of the two perpendicular arms, a small metal bowl contained 1/8th of a chocoloop (Kellogs) for the rewarded phase of the behavioural protocol. Visual cues were disposed around the maze for spatial orientation. Apparatus was in an isolated room with dim lighting (5-6 lux) to avoid stressing the animals.

Animals were habituated to the experimental procedure over 2 days. On the first day, mice were placed in groups (5-6 mice) in the T-maze to freely explore the maze for 10 min. The floor of the maze was covered with chocoloop (Kellogs) as a reward to encourage the animals to enter the T-maze. The second day was identical to the first day, but animals were placed individually in the maze for the 10 min exploratory phase and food was only at the reward place in the metal bowl at the end of both arms. From the habituation onwards, mice were gradually food restricted to 85% of their body weight. To implement this restriction, the animals’ weights were monitored daily through weighing. Following each weighing, the quantity of food was adjusted to maintain their body weight at 85%. The food was then distributed directly to their housing cages.

For DNMP training (Figure 1A), each trial was divided into two phases separated by a delay. During the sample phase, one of the two arms was blocked, forcing the mouse to move to the arm that remained open (left or right). During the choice phase, both arms were open and the mouse had to choose to move to either arm. The trial was considered correct if the mouse visited the previously closed arm. If the mouse made the correct choice, it received a food reward (chocoloop). The delay between the two phases was set at 90 seconds for the whole duration of the experiment. These two phases were repeated 10 times (10 trials) per day per mouse. Within a trial, the blocked arm during the sample phase was decided randomly and changed between trials. This sequence of 10 trials was itself randomly determined and changed each day according to specific criteria: no more than 3 consecutive trials per day with the sample in the same arm and every day each animal had always the same number of samples left and right. For the full protocol, an animal was considered to have reached the success criterion when it obtained an average of 8 out of 10 successful attempts (80% success rate) during three consecutive days. Based on this criterion the task was separated into two parts. The first part from the beginning of the experiment until the animals reached the success criterion, was called the learning phase. The second part, once the mice have reached the success criterion, was called the maintenance phase of memory. Some batches of mice were trained until reaching a different sub-optimal criterion of 70% success rate during 2 non-consecutive days to test their behaviour in the devaluation protocol during the learning phase.

**Figure 1.**
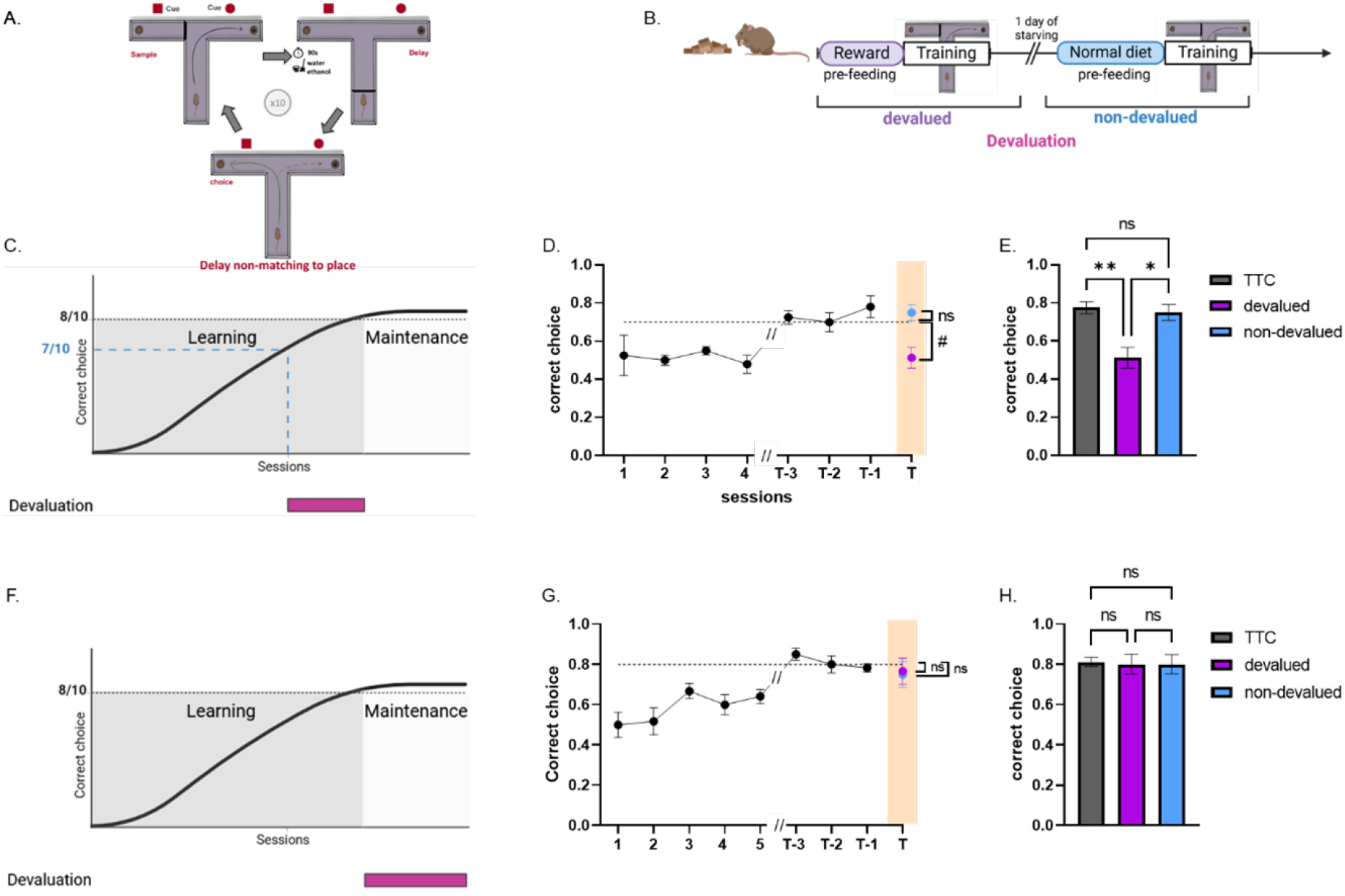
Shift of strategy from goal-directed to automated behaviour in the DNMP task. **A**. Diagram of a DNMP session in the T-maze. **B**. Devaluation protocol in the DNMP task. The value of the reward was decreased by pre-feeding with reward food delivered in the T-maze for one-hour before training. Satiety effect on behaviour in the T-maze was tested by pre-feeding with a normal diet (cage food). **C**. Experimental design of devaluation time during the learning phase of DNMP task. **D**. Learning curve per session until the 70% criterion was reached during 2 non-consecutive days (black dots) and after devaluation and success score after devaluation (purple dot) or non-devaluation (blue dot) protocols. **E**. Bar graph of correct choices in DNMP task upon reaching criterion (Time to criterion (TTC): average of last three days of training before devaluation), after devaluation or non-devaluation protocols (n=8). **F**. Experimental design of devaluation time during the maintenance phase of DNMP task. **G**. Learning curve per session until the 80% criterion for 3 consecutive days was reached (black dots) and success score after devaluation (purple dot) or non-devaluation (blue dot) tests. **H**. Bar graph of correct choices in DNMP task upon reaching criterion (TTC), after devaluation or non-devaluation protocols (n = 12). Statistics: D: One sample t test, devalued x hypothetical value (0.7), non-devalued x hypothetical value (0.7), #p=0,0112, “ns” p = 0,2753; E: One-way Anova, TTC x devalued x non-devalued, **p=0,0028, “ns” p = 0,6682, *p=0.02; G : One sample Wilcoxon test, devalued x hypothetical value (0.8), non-devalued x hypothetical value (0.8), “ns” p=>0,9999/ >0,9999; H : Friedman test, TTC x devalued x non-devalued, “ns” p= >0,9999 / >0,9999 / >0,9999. Error bars represent SEM. Drawings created with BioRender.com. Full statistics can be found in Supplementary Statistics Table Figure 1.

### Devaluation protocol in T-maze

The devaluation procedure is presented in Figure 1B. Two criteria based on the T-maze success score were set up to separate two groups subject to devaluation. In this way, one group of mice was devalued during the learning phase and one group during the maintenance phase. For the first group to be devalued during learning, animals had to achieve a 70% success rate during 2 non-consecutive days before being devalued. For the second group devalued during the maintenance of phase, animals had to first achieve the success criterion used in the DNMP, i.e. 80% success rate, before devaluation. Each of the two groups was then subjected to a two-day devaluation extinction test of a specific type of reward, during which the behaviour in the T-maze for the DNMP was measured. On the day following the last day of training, all animals were given free access to either reward food (chocoloops) or usual food diet for one hour. Immediately after this pre-feeding treatment, animals were trained in the T-maze for a 10-trial devaluation session in which correct choices in the DNMP were measured in the absence of reward. A second test was conducted the following day. It was identical to the first day, but the animals were fed with the second type of food, either usual food or reward food depending on what they had been exposed to on the first day. The two successive devaluations were separated by 24 hours of starving to reverse the effect of the first devaluation before the second devaluation. According to this protocol, mice were divided into two groups: the devalued group, which was pre-fed with the food reward, and the non-devalued group, which was pre-fed with their regular food.

### Locomotor activity

An open-field apparatus was used to measure locomotor activity and stress. Each mouse was placed in a 40 × 40 cm square open field with an ambient light of 200 lux for 5 min and left to explore freely. Mice movements were recorded using a video camera placed above the apparatus. Distance travelled, times in central zone and peripheries zone were automatically analyzed using the software AnyMaze (Stoelting, France).

### Anxiety test

O-maze apparatus was used to assess anxiety levels. O-maze test was conducted in a white circular runway (55 cm diameter, ring shaped) raised 60 cm above the floor. The corridor width was 5 cm and was divided into two open quadrants (open zone) facing two enclosed quadrants (close zone) with a 15 cm high wall. Mice were placed in an open zone facing a closed zone and allowed to explore for 10 min with an ambient light of 200 lux. Time spent in open zones and close zones was automatically analyzed using the software AnyMaze (Stoelting, France).

### Statistical analysis

Data were analyzed using Graph Prism 8.0 (GraphPad, U.S.A.). Normality of the distribution was first tested using a Kolmogorov–Smirnov test and D’Agostino & Pearson test. Depending on the number of groups and the results of the normality test, groups were then compared using a One-sample t test, Unpaired t test, Paired t test, Wilcoxon test, one-way or two-way analyses of variance followed by post hoc Sidak’s test. Data in the figures are presented as mean ± SEM. Statistical significance was set at P < 0.05. Statistical tables for figure 1 to 6 are reported in supplementary Table 1 to 6.

## RESULTS

### Behavioural transition from goal-directed to automated behaviour during the evolution of working memory

Our goal was to analyze if different types of behaviours (eg. goal-directed or automated) were at play during the different phases of working memory processing. We therefore asked if the behavioural strategy of mice changed during a DNMP (Figure 1A). For this purpose, together with the DNMP task, we adapted a devaluation task commonly used in operative cages to analyze if the behaviour was goal-directed or automated (Killcross, 2003) to the T-maze (Figure 1B). For devaluation, we carried out a one-hour pre-feeding with either the food that the animal normally received in the cage or with the food used as a reward in the T-maze. After a 24-hour starving period, the animals were given a second devaluation with the other type of food. We performed the devaluation two times during the DNMP task: either during the learning phase or during the maintenance phase (Figure 1C and F). The moment of devaluation during the learning phase was set at 7 out of 10 correct trials for two non-consecutive days (Figure 1C). We observed that, during the learning phase, submission of mice to the devaluation protocol was associated with a drop in the success score in the T-maze task (Figure 1D and 1E). This alteration was not due to the effect of satiety alone associated with pre-feeding, as non-devaluated mice (still fed to satiety with normal diet) did not exhibit altered memory (Figure 1D and 1E). These data demonstrate that mice exhibit goal-directed behaviour that is sensitive to the value of the reward during the learning phase of this working memory task.

To evaluate the type of behavioural strategy used by the mice during the maintenance phase of this working memory task, we performed the devaluation protocol after mice had reached a more stringent DNMP criterion (80% correct choices for 3 consecutive days) (Figure 1F). In this maintenance phase, the devaluation of the reward did not reproduce the effects observed during the learning phase, in terms of the success score in the DNMP (Figure 1G and 1H). Indeed, devaluated mice still exhibited a strong success score identical to non-devaluated mice and to the success score observed before devaluation. Our results demonstrate that, once the mice have reached the maintenance phase of the memory, mice adopt automated behaviour that is less sensitive to the value of the reward. Together, these results argue for a change in strategy during the DNMP task, where devaluation-sensitive goal-directed behaviour is lost upon robust encoding of this working memory, with a shift towards automated behaviour.

### Role of the mPFC during working memory

The change in behavioural strategy observed during DNMP could be based on specific brain dynamics. Our hypothesis was that the mPFC would be important during the learning phase, where goal-directed behaviour is evident (Figure 1D and 1E) (Delatour and Gisquet-Verrier, 2000; Ragozzino et al., 1999), while other structures, such as the striatum, might contribute to the automation of the memory during the maintenance phase (Figure 1G and 1H) (Coutureau and Parkes, 2018). To test this hypothesis, we used chemogenetics to inhibit neuronal activity in the mPFC during the different phases of the DNMP task.

Viral expression was targeted to the mPFC. Figure 2A displays a typical expression of hM4Di together with mCherry in the mPFC. We inhibited neuronal firing by CNO or saline i.p. injection every day 30 minutes before the start of the training until one of the two groups reached the success criterion (Figure 2B). As expected, with this protocol, control (saline-injected) mice reached the stringent 80% correct choices for 3 consecutive days, while mPFC-inhibited mice failed to reach this criterion (Figure 2C). This result is confirmed when the average score of correct choices of the last three days of training is represented (Figure 2D). These results allow us to confirm the necessity of mPFC neuronal activity during the learning phase of the DNMP task. We then tested the involvement of the mPFC during memory maintenance. For this, we performed inhibition of the mPFC by injection of CNO for two consecutive days only after all mice had reached the stringent criterion of 80% correct choices for 3 consecutive days (Figure 2E). To allow animals to act as their own controls, half of these mice received CNO on the first day (30 min before performing the task) and the other half on the second day allowing for a 24h period between the two days to ensure full elimination of CNO. Inhibition of neuronal activity in mPFC during the maintenance phase did not have any effect on the performance in the DNMP task as measured by the success score of correct choices (Figure 2F and 2G). These data show that inhibition of the mPFC impacts the behaviour only during the learning phase, but not during the maintenance phase of the DNMP task. These data suggest that other structures are implicated during maintenance of this memory, leading to an automation of the task.

**Figure 2.**
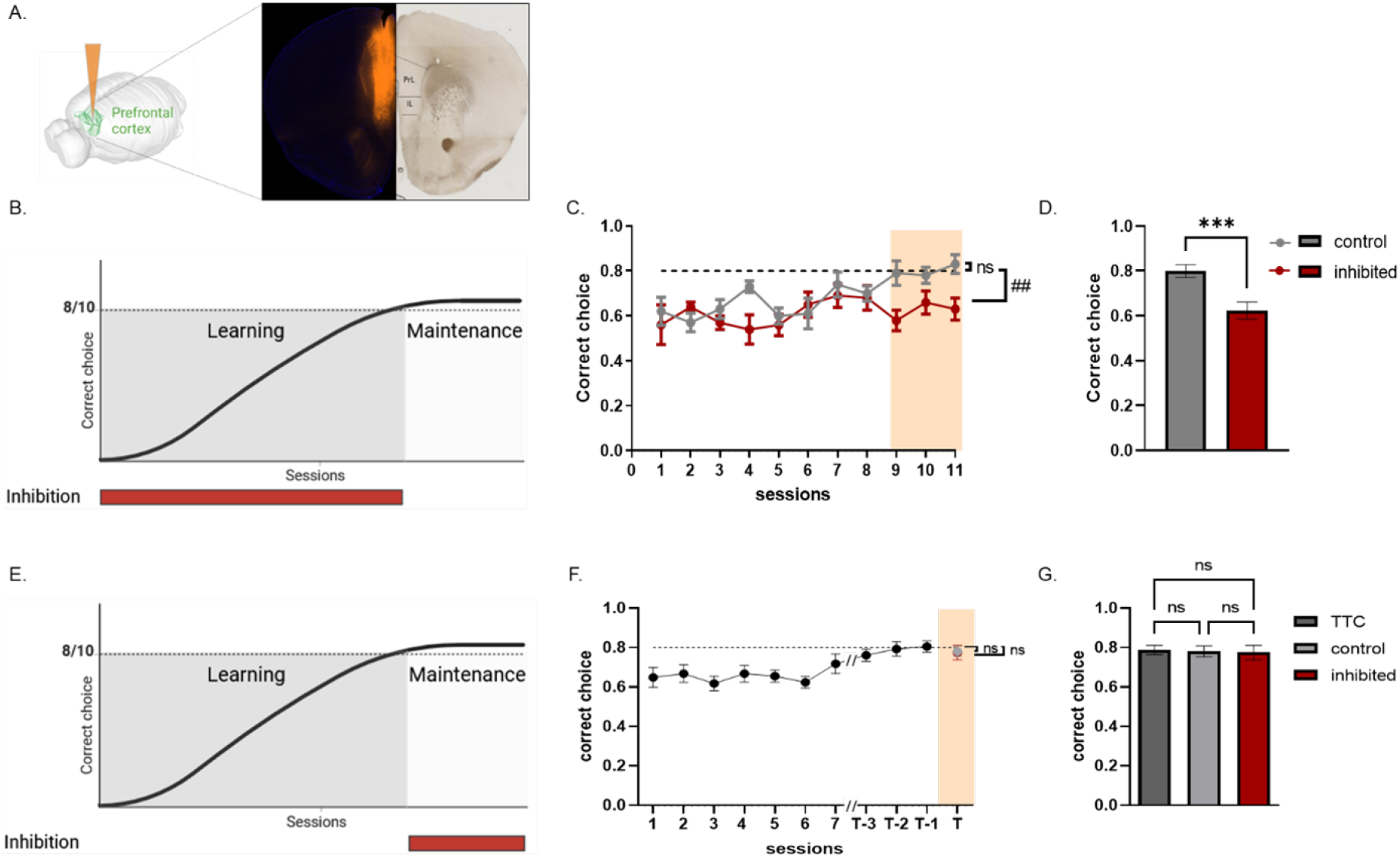
The mPFC is necessary during the learning phase but not the maintenance phase of the DNMP task. **A**. Example of hM4Di in vivo viral expression with mCherry fluorescence (orange) in the mPFC. **B**. Experimental design with time of mPFC-specific inhibition of neurons during the learning phase of DNMP task. **C**. Learning curve per session of mPFC-inhibited (CNO; red; n=10) and control (Saline; grey; n=10) mice per session. **D**. Bar graph of correct choices in DNMP task (average of last three days) for mPFC-inhibited and saline-injected control mice. **E**. Experimental design of time of mPFC-specific inhibition of neurons during the maintenance phase of DNMP task. **F**. Learning curve per session before mice reach the success criterion (black dots) and results for mPFC-inhibited mice (CNO; red) and saline-injected control mice (Saline; grey) after criterion is reached. (n=16). **G**. Bar graph of correct choices in DNMP task for criterion (TTC: average of the last three days of training before injection), mPFC-inhibited and saline-injected control mice. Statistics: C: One-sample t test, sessions 9 / 10 / 11: mean control x hypothetical value (0.8), mean inhibited x hypothetical value (0.8), “ns” p= >0.9999, ##p= 0,0012; D: Unpaired t test, mean 3 last days control x means 3 last days inhibited, ***p=0.0008. F: One sample t test, control x hypothetical value (0.8), inhibited x hypothetical value (0.8), “ns” p= 0,5104/ 0,5090; G: One-way Anova, criterion x control x inhibited, “ns” p= 0,9779/ 0,9779/ 0,9779; Error bars represent SEM. Full statistics can be found in Supplementary Statistics Table Figure 2. Drawings created with BioRender.com.

### Role of the dorsolateral striatum during working memory

As we observed an automation of the behaviour during the maintenance phase of the memory in the DNMP task, we set out to determine the involvement of the DLS, one of the structures most associated with this type of behaviour (Coutureau and Parkes, 2018). As done for the mPFC, we inhibited neuronal activity in the DLS by chemogenetics either throughout the learning phase (Figure 3A-B) or just during the maintenance phase (Figure 3F), hypothesizing that DLS might be playing a role only during the maintenance phase. Figure 3A displays a typical expression of hM4Di together with mCherry in the DLS. We performed CNO injections as for the mPFC experiments (30 minutes before task). When DLS inhibition occurred throughout the learning phase, this did not hinder learning of the task as mice reached the success criterion as did control saline-injected mice (Figures 3C-D). When we inhibited neuronal activity in the DLS with CNO only during the maintenance phase, once the animals had reached the stringent success criterion (80% of correct choices for 3 consecutive days), we also observed normal maintenance in DLS-inhibited mice (Figure 3G and H). This suggests that the DLS is not necessary to encode or maintain this memory.

**Figure 3.**
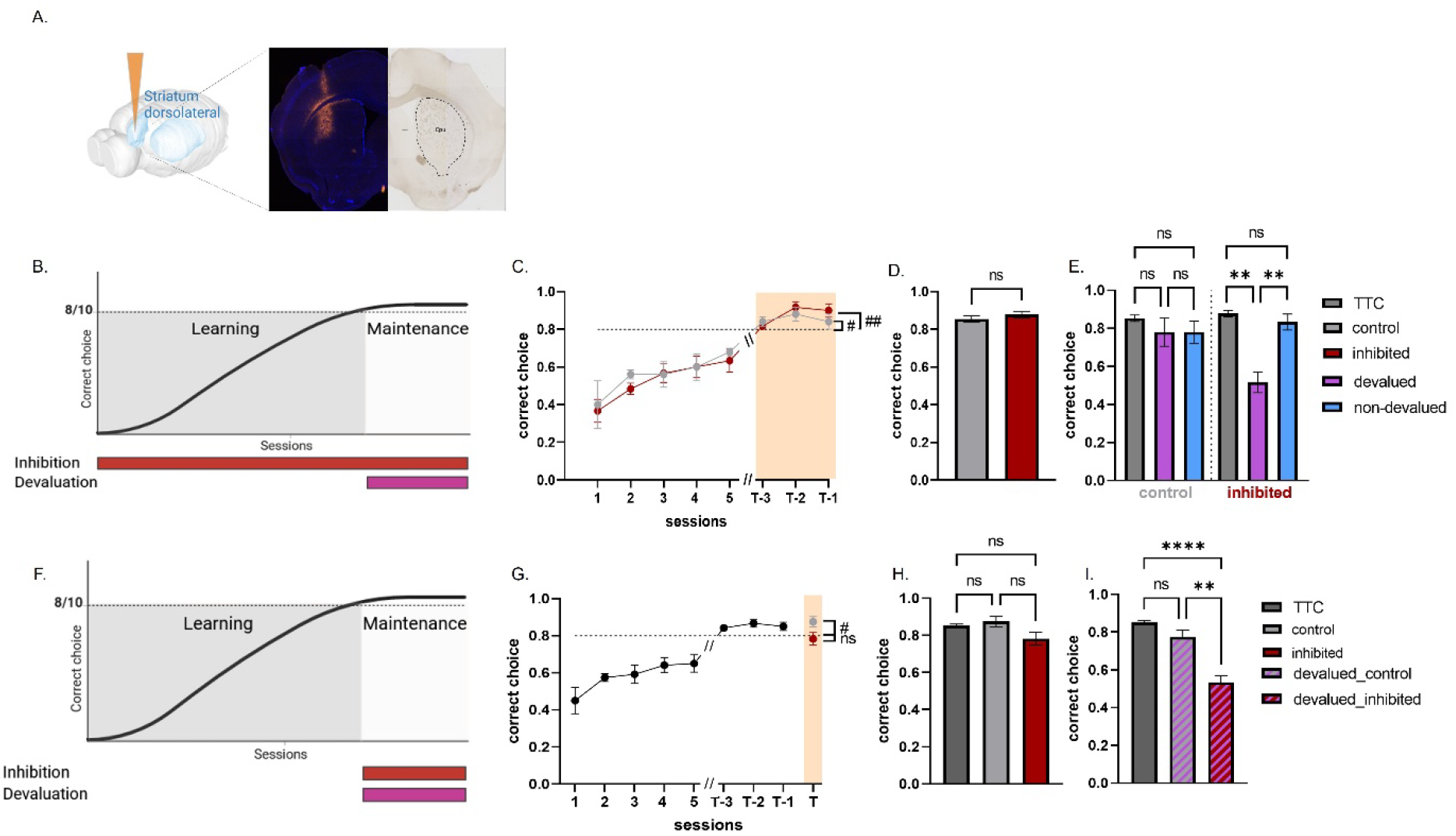
The DLS is necessary for transition from goal-directed to automated behaviour during the DNMP task. **A**. Visualization of hM4Di *in vivo* viral expression with mCherry fluorescence (orange) in the DLS to obtain DSL-specific inhibition of neuronal activity. **B**. Experimental design of timing of DLS inhibition and devaluation during learning phase of DNMP task. **C**. Learning curve per session of DLS-inhibited mice (CNO; n=6) and control mice (Saline; n=5). **D**. Bar graph of correct choices made by DLS-inhibited and saline-injected control mice upon mice reaching criterion. **E**. Bar graphs of correct choices of control and DLS-inhibited mice upon reaching criterion (TTC, average of the last three days of training before devaluation), in devalued and non-devalued mice. **F**. Experimental design of timing of DLS inhibition and devaluation during the maintenance phase of DNMP task. G. Learning curve per session of DLS-inhibited and control mice to reach criterion (n=12). H. Bar graphs of correct choices for mice once they reach the criterion (TTC) and after DLS-inhibition (red dot) or control saline (grey dot) injection. I. Bar graph of correct choices reached by mice upon reaching criterion compared to DLS-inhibited or saline-injected control mice after devaluation protocol. Statistics: C: One sample t test, session T-3/T-2/T-1, mean control x hypothetical value (0.8), mean inhibited x hypothetical value (0.8), #p=0.0349, ##p=0.0052; D: Unpaired t test, mean 3 last days control x means 3 last days inhibited, “ns” p=0.3316 ; E : One-way Anova, control :TTC x devalued x non-devalued, “ns” p= 0.5659 / 0.3054 / >0.9999; One-way Anova, inhibited : TTC x devalued x non-devalued,**p= 0.0027 / 0.0094, “ns” p=0.3075; G : One sample t test, control x hypothetical value (0.8), inhibited x hypothetical value (0.8), #p=0.0210, “ns” p=0.6380, H : One-way Anova, TTC x control x inhibited, “ns” p= 0.4524 / 0.1736 / 0.1429; I : One-way Anova, control : criterion x devalued-control x devalued-inhibited, **** p=<0.0001, ** p=0.0031. Error bars represent SEM. Full statistics can be found in Supplementary Statistics Table Figure 3. Created with BioRender.com.

In both these control and DSL-inhibited mice, we however also interrogated the strategy employed during the task by submitting them to the devaluation protocol during the maintenance phase, once the stringent criterion was reached. As expected, in the control saline-injected mice, devaluation had no effect on the correct choices obtained in the DNMP task (Figure 3E and I), showing a shift towards automated behaviour as previously evidenced in Figure 1H. However, when DLS was inhibited throughout learning or just during maintenance (Figure 3B and F), devaluation had a significant effect on this task (Figure 3E and I). These data suggest that, due to the absence of activity of the DLS, mice resorted back to a goal-directed strategy for maintenance.

Together, these results show that, although DLS is not necessary to perform this task, its inhibition prevents the transition from a goal-directed to an automated behaviour.

### mPFC and DLS are both necessary during the maintenance phase of working memory

Considering that DLS-inhibited mice could shift back to a goal-directed strategy when DLS was inhibited, we hypothesized that, although the mPFC is not necessary to perform the task during the maintenance phase (Figure 2F-G), it might not be completely disengaged and could compensate for this maintenance if necessary. We expressed hM4Di in both the mPFC and DLS to inhibit both structures during the maintenance phase (Figure 4A and 4B). Again, for this experiment, we performed inhibition of both structures by injection of CNO or saline for two consecutive days only after all mice had reached the stringent criterion of 80% correct choices for 3 consecutive days (Figure 4B-C). To allow animals to act as their own controls, half of these mice received CNO on the first day (30 min before performing the task) and the other half on the second day allowing for a 24h period between the two days to ensure full elimination of CNO. As hypothesized, when both structures were inhibited, mice did not maintain the memory adequately when compared to saline-injected control mice (Figure 4C-D). No alterations in motor activity or anxiety-like behaviour were observed in these mPFC/DLS-inhibited mice (supplementary Figures S1 and S2). These data argue that the mPFC is not fully disengaged for the maintenance of working memory in this task and can be re-engaged when the DLS is not available, explaining probably why DLS-inhibited mice could still switch back to a goal-oriented behaviour.

**Figure 4.**
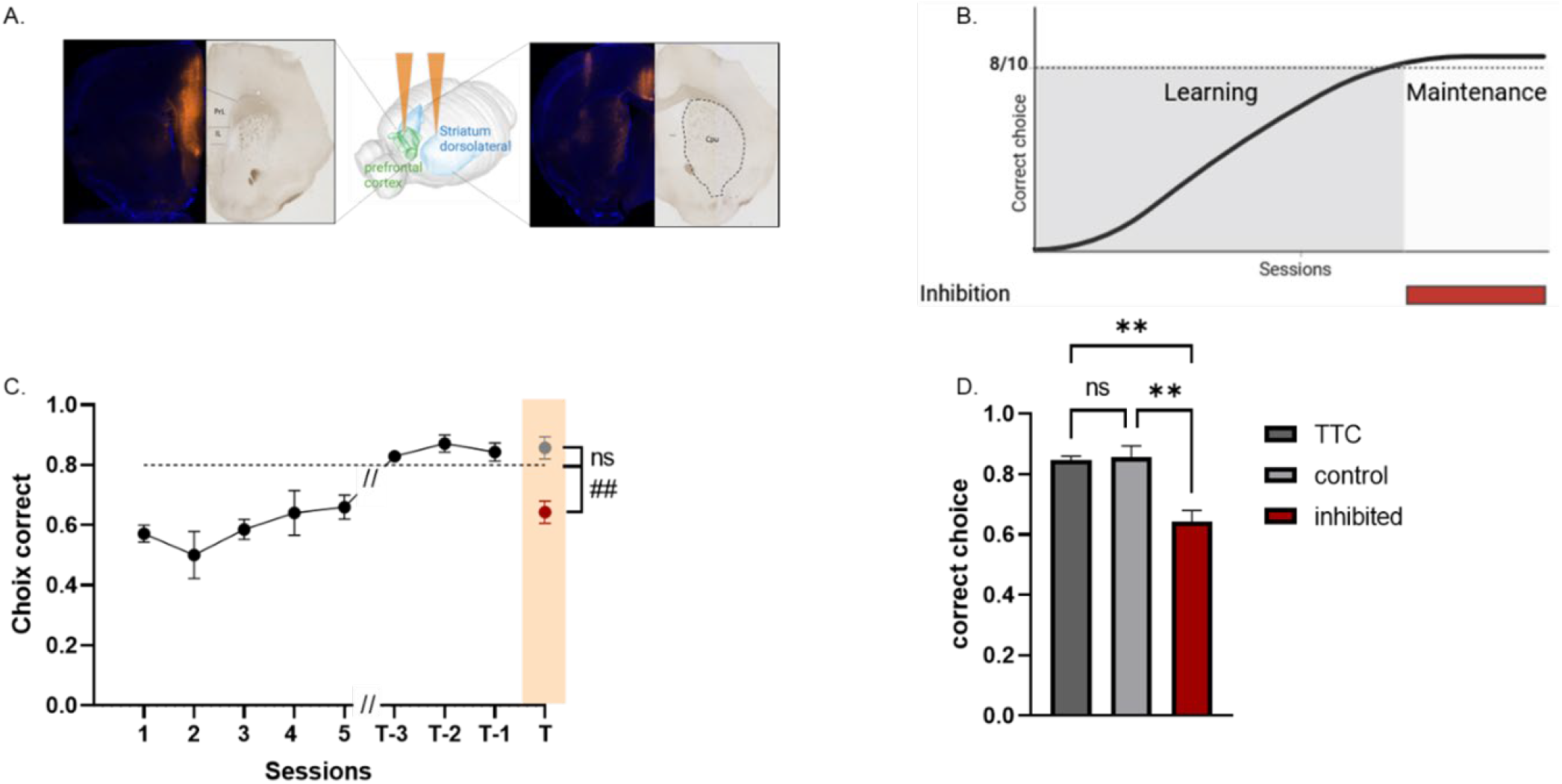
Memory maintenance in DNMP task is disrupted when neuronal activity is inhibited in both mPFC and DLS. **A**. Visualization of hM4Di in vivo viral expression with mCherry fluorescence (orange) in the mPFC and DLS. **B**. Experimental design of inhibition during the maintenance phase of DNMP task. **C**. Learning curve of correct choices of mice once they reached the criterion and in mPFC/DLS-inhibited (CNO; red dot) and control (Saline; grey dot) mice (n=7). **D**. Bar graphs of correct choices for criterion (TTC, average of the last three days of training before injection), mPFC/DLS-inhibited and saline-injected control mice. Statistics: C: One sample t test, control x hypothetical value (0.8), inhibited x hypothetical value (0.8), “ns” p= 0.1723, ##p= 0.0053; D: One-way Anova, TTC x control x inhibited, “ns” p= 0.8119, **p= 0.0078, **p= 0.0078. Error bars represent SEM. Full statistics can be found in Supplementary Statistics Table Figure 4. Drawings created with BioRender.com.

## DISCUSSION

Our study assessed the involvement of the mPFC and DLS during the learning and maintenance phases of a spatial working memory task. In particular, we questioned the capacity for automation of this task. We show here that the mPFC, but not the DLS, was essential for learning the task. This result was expected from the well-known importance of the mPFC in the encoding of working memory (Funahashi, 2017). However, by contrast, inhibition of the mPFC during the maintenance phase had no effect on memory performance, suggesting that another brain structure may take the relay. This shift strongly suggests the potential involvement of structures involved in procedural memory, such as the DLS. Yet, surprisingly, inhibiting the DLS during maintenance had no effect on memory performance. Interestingly, results of the devaluation protocol performed on these DLS-inhibited mice demonstrated that these mice shifted their behaviour back to a goal directed strategy. Finally, when both the mPFC and DLS were inhibited concomitantly during the maintenance phase, mice did not manage to maintain this memory, arguing for a necessary interplay between the two structures. More specifically, we demonstrate that automation of this type of memory during the maintenance phase is possible with a potential involvement of the DLS, but that the mPFC is not fully disengaged. This suggests that the memory engram in the mPFC may remain in a dormant state, with the potential to be reactivated whenever necessary.

### Goal directed versus automation strategy

We adapted the devaluation protocol commonly used in operant cages to assess different behavioural strategies (Coutureau and Parkes, 2018; Yin et al., 2005, 2004) for the DNMP task. The value of the reward was reduced by pre-feeding the animals with the food they would receive as a reward before submitting them to another trial of the DNMP task. Consequently, the decrease in the DNMP score with devaluation (with reward) during learning indicated that the animals adopted a goal-directed behaviour. In contrast, the lack of effect of devaluation on mice during the maintenance phase of this memory suggested that the mice had adopted a more rigid automatized behaviour insensitive to changes in reward value(Killcross, 2003). These results indicate a transition from this goal-directed behaviour to automatized behaviour after prolonged training. The emergence of this rigid behaviour can be explained by the characteristics of our DNMP task, which allows automation of the rule. Unlike previous studies, in our protocol, we used a fix delay between sample and test and a long training due to high success criterion. Consequently, we argue that the stability of the task reinforces the animal’s integration of the procedure. However, these training characteristics do not eliminate the use of working memory by the animals. Despite the invariability of most of the parameters of our task (maze, cues, delay, rule), the sequence of the ten daily sessions (direction of the forced arm linked to the direction of the choice arm) was determined randomly and changed each day, thus preventing the animal from retaining a specific sequence. The task, therefore, required the forced arm information to be updated between each of the ten sessions to prevent previous information from disrupting the choice in the following session.

### Brain structures associated to goal-directed strategy and automation strategy

Here, with the use of chemogenetics, we also analyzed which brain structures might be responsible for the evolution of strategies in this working memory. We began by conditional inactivation of the mPFC either during the learning phase or during the maintenance phase of the DNMP task. Inhibition of the mPFC during learning prevented animals from reaching the success criterion, highlighting the previously evidenced crucial role of this structure in learning of a working memory task (Euston et al., 2012; Vogel et al., 2022). Conversely, inhibition during the maintenance phase had no impact on memory performance. It appears that the mPFC, essential during the learning phase when behaviour is goal-oriented, is disengaged during the maintenance phase when behaviour becomes automated. Similar results were reported by Tunes et al. (Tunes et al., 2022) where, in a simple motor timing task, the authors found that mPFC inactivation severely impaired the learning phase but had no effect on the maintenance phase. Our results do not exclude the possible role of the mPFC during the maintenance phase, which always engages working memory (Vogel et al., 2022), but its role can be compensated if this mPFC is dysfunctional —compensation that would be impossible when mPFC damage occurs during the learning phase. This result supports the idea that task-induced overtraining favors automated strategies, allowing other structures, such as the striatum, to take over and rendering the mPFC non-essential to the task (Hasz and Redish, 2018).

This idea is supported by our results, which show that inhibition of the DLS during the maintenance phase prevents the formation of automated, devaluation-insensitive behaviour, whereas it has no effect when this inhibition is during the learning phase. Involvement of the DLS in automated behaviour is well-recognized in the literature (Balleine and Dickinson, 1998; Balleine and O’Doherty, 2010; Dickinson, 1994; Dickinson et al., 1995; Graybiel, 1998; Hilario et al., 2012b; Yin et al., 2004, 2005). Neuronal activity in the DLS has also been associated with a task-bracketing pattern, marking the beginning and end of action sequences in well-learned behaviours, and this activity is insensitive to devaluation (Smith and Graybiel, 2013) such as in our results. The fixed 90-second interval between the sample and test phases in our experiment likely reinforced the task-bracketing effect, which may explain the prominent role of the DLS in our task. In a more recent study, decoding analysis of neuronal activity revealed a shift in the control of temporal responses (i.e., time intervals between the sample and test phases) from the mPFC during the early stages of learning to the DLS during the maintenance phase in a simple motor task (Tunes et al., 2022). This study is in accordance with our results but we can now be extrapolated to more complex tasks such as working memory tasks.

Finally, our last experiment tackled a very interesting concept. Indeed, in our experiment, inhibition of the DLS did not directly affect the animal’s performance during learning and maintenance. Interestingly, DLS inhibition during the maintenance re-engaged a goal directed behaviour that is supported by the mPFC. This relay was evident as memory maintenance was lost when both structures were inhibited simultaneously. This experiment suggests that the memory trace within the mPFC network is neither erased nor transferred to the dorsal striatum, but rather remains in a dormant or silent state—, ready to be re-engaged when cognitive flexibility is required as suggested previously (Tonegawa et al., 2018)). The question that arises, then, is: why are different strategies supported by distinct neural networks? The most plausible answer is a matter of cost-benefit trade-off, as developed in many reinforcement models (James et al., 2023).

This adaptation of behavioral strategies may reflect a reduction in memory load, shifting from a cortical flexible system, involved in many high cognitive functions and highly connected to many structures, such as the mPFC, to a more subcortical automated system, involved in inflexible routine, such as the DLS.

Our results support the possible coexistence of several behavioural strategies, with the most adapted, cost-effective strategy expressing itself naturally as suggested by recently of (Bouton, 2021).

## Acknowledgment

This work was supported by the French government through the France 2030 investment plan managed by the National Research Agency (ANR), as part of the Initiative of Excellence Université Côte d’Azur under reference number ANR-15-IDEX-01, and in particular by the interdisciplinary Institute for Modeling in Neuroscience and Cognition (NeuroMod) of the University Côte d’Azur (Villet PhD fellowship) and by CNRS : MITI notification Défi - Modélisation du vivant 2018-2019 Projet : DYNA MO -BETHUS Ingrid.

## SUPPLEMENTARY FIGURES AND TABLES

**Figure S1.**
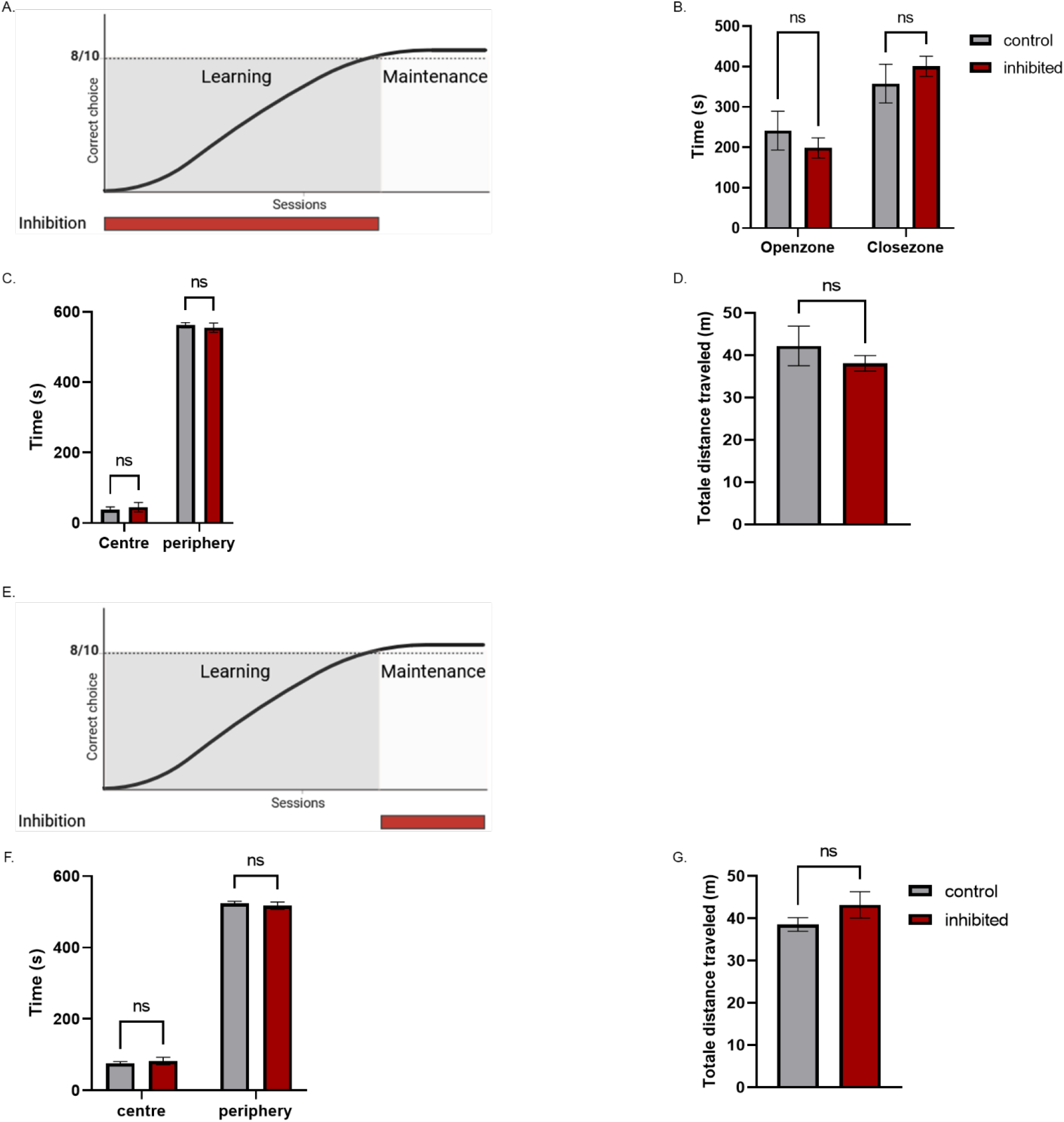
Evaluation of locomotion and anxiety in mice with inhibition of the DLS. **A**. Experimental design of inhibition during the learning phase of DNMP task. **B**. Average time spent by animals in the open and closed areas of the O-maze during ten minutes. **C**. Average time spent by animals in the central and peripheral zones of the open field during ten minutes. **D**. Total distance covered by animals in an open field. **E**. Experimental design of inhibition during the maintenance phase of DNMP task. **F**. Average time spent by animals in the central and peripheral zones of the open field during ten minutes. **G**. Total distance covered by animals in an open field. (control n= 5, inhibited n= 6; B: Two-way anova, open zone: control x inhibited, “ns” p=0,6535, close zone: control x inhibited, “ns” p= 0,6535; C: Two-way anova, centre: control x inhibited, “ns” p= 0,8873, periphery: control x inhibited, “ns” p= 0,8873; D: Unpaired t test, control x inhibited, “ns” p= 0,4028; control n= 7, inhibited n= 7; F: Two-way anova, centre: control x inhibited, “ns” p= 0,8007, periphery: control x inhibited, “ns” p= 0,8007; G : Paired t test, control x inhibited, “ns” p= 0,2463).

**Figure S2.**
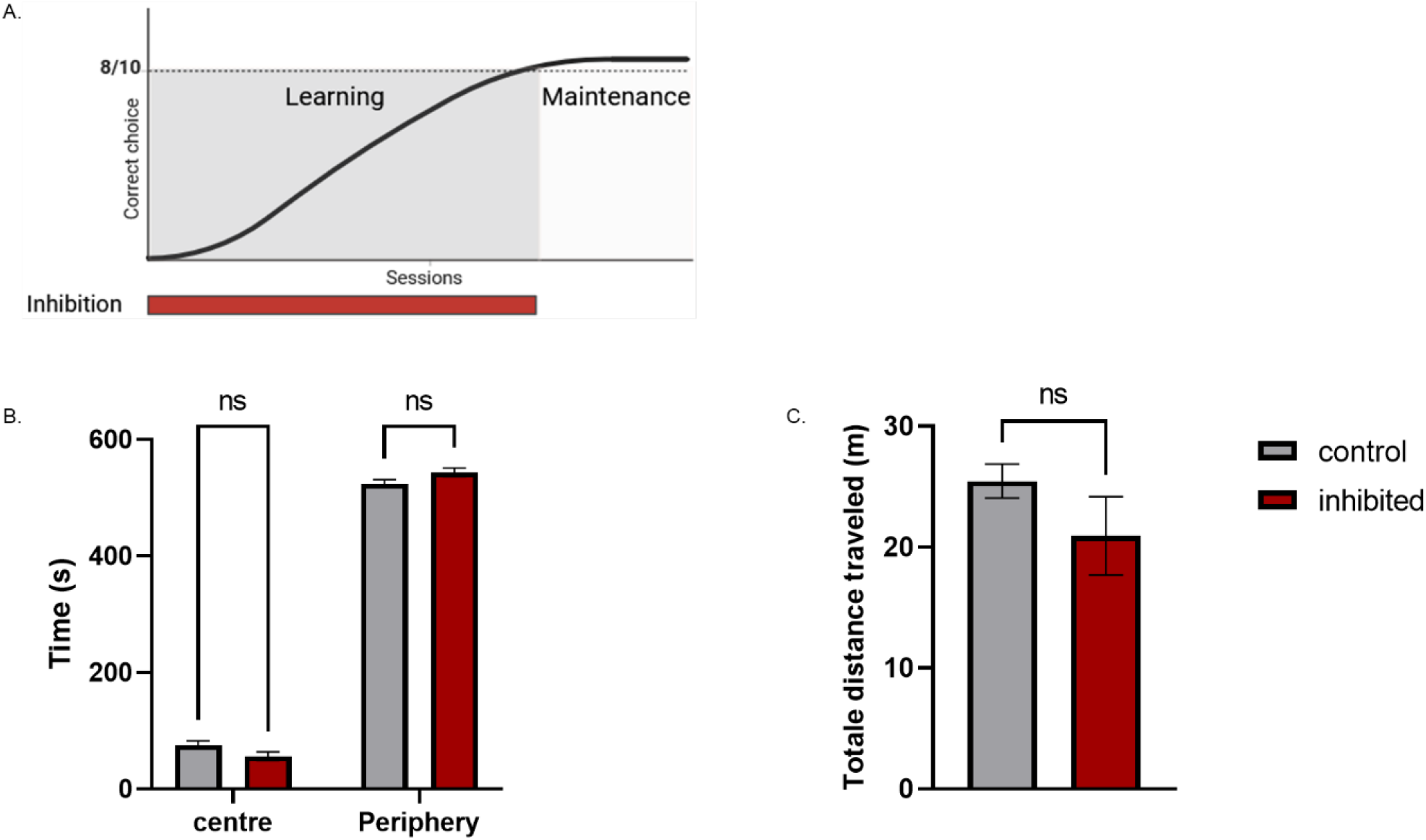
Evaluation of locomotion and anxiety in mice with inhibition of mPFC and the DLS. **A**. Experimental design of inhibition during the maintenance phase of DNMP task. **B**. Average time spent by animals in the central and peripheral zones of the open field during ten minutes. **C**. Total distance covered by animals in an open field. (control n= 5, inhibited n= 4; B: Two-way anova, centre: control x inhibited, “ns” p= 0,1545, periphery: control x inhibited, “ns” p= 0,1545; C: Mann-Whitney test, control x inhibited, “ns” p= 0,4127).

**Table S1.**
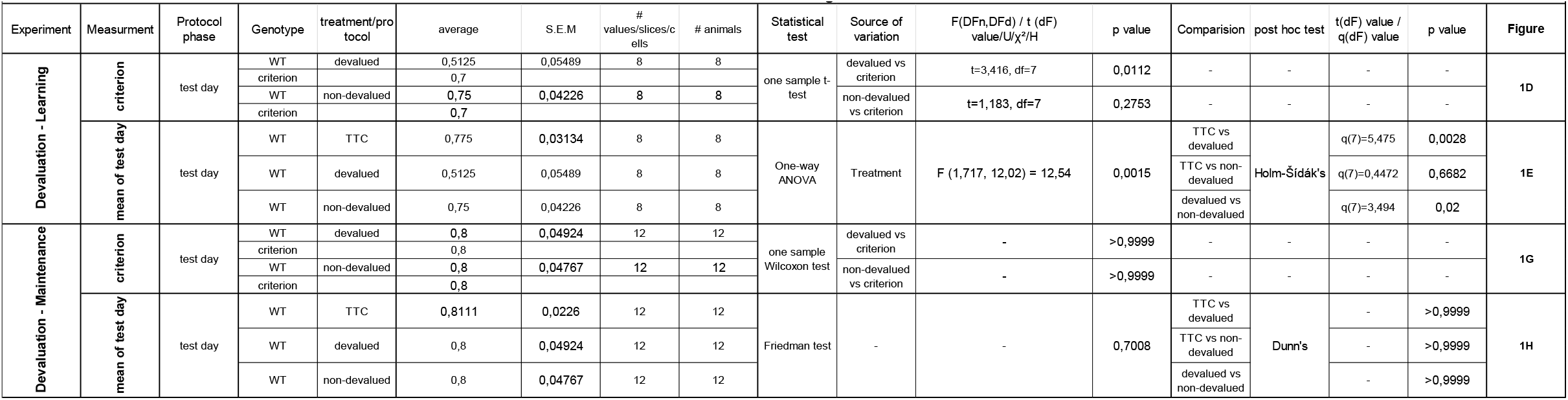
Statistics Table Figure 1.

**Table S2.**
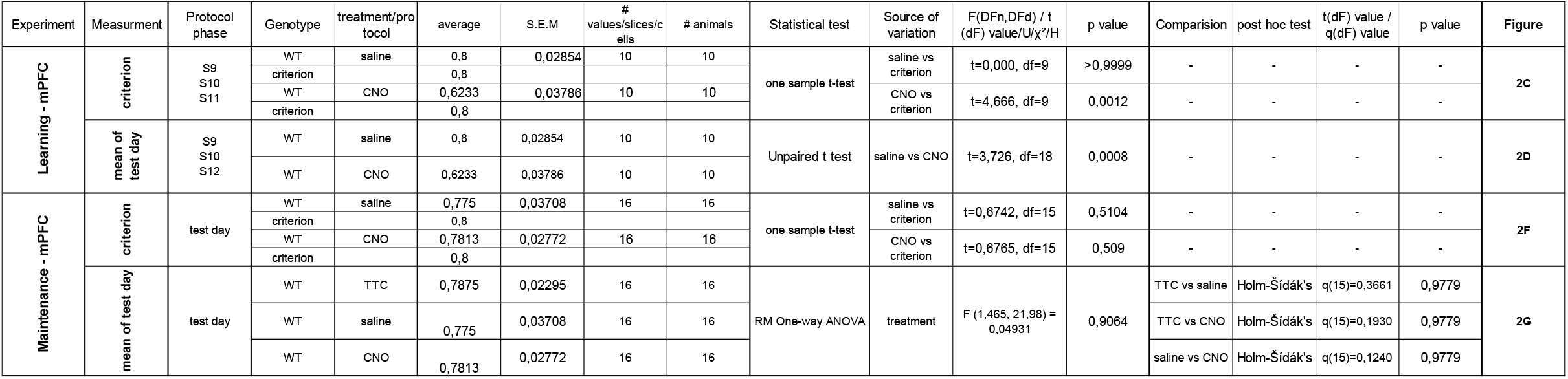
Statistics Table Figure 2.

**Table S3.**
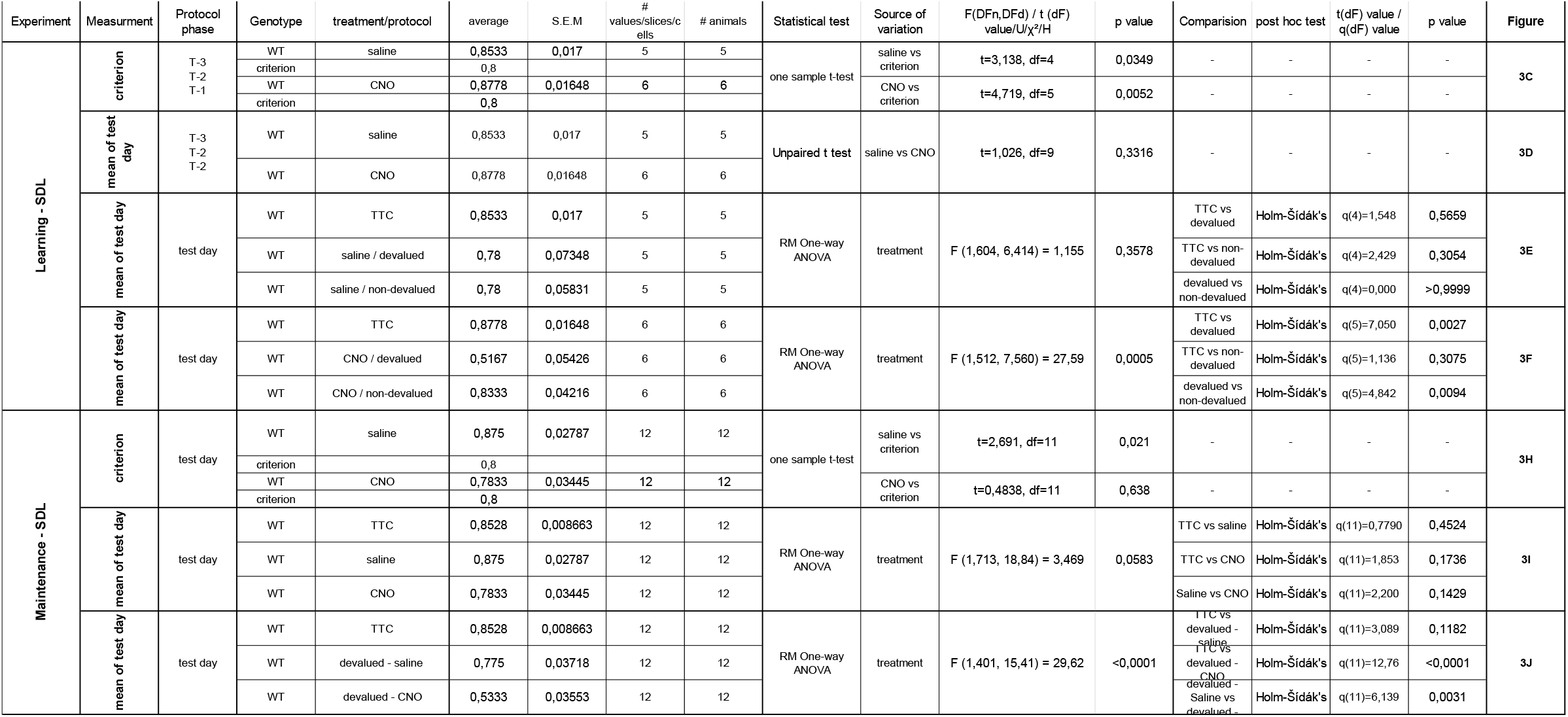
Statistics Table Figure 3.

**Table S4.**
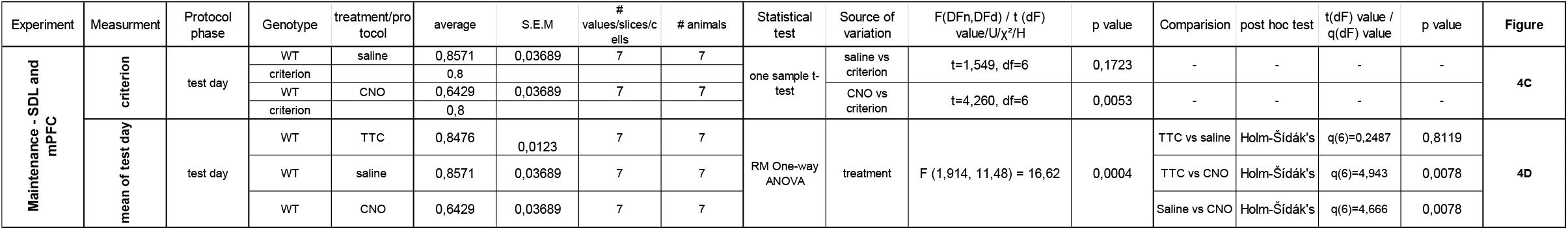
Statistics Table Figure 4.

**Table S5.**
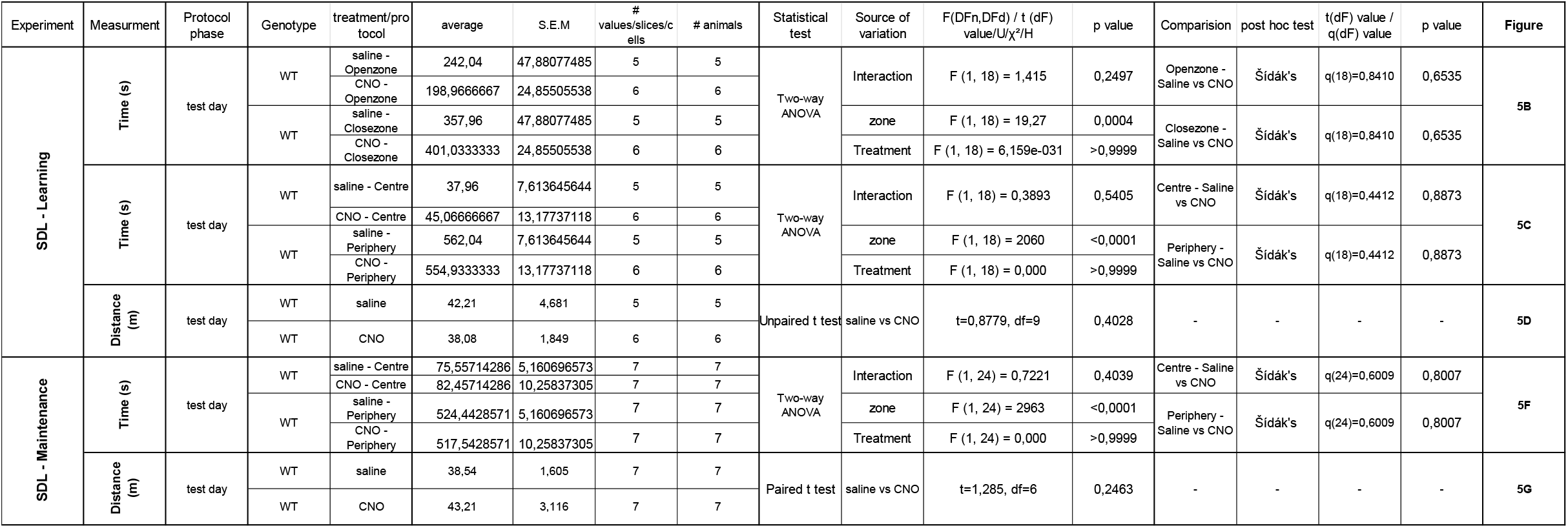
Statistics Table Figure 5.

**Table S6.**
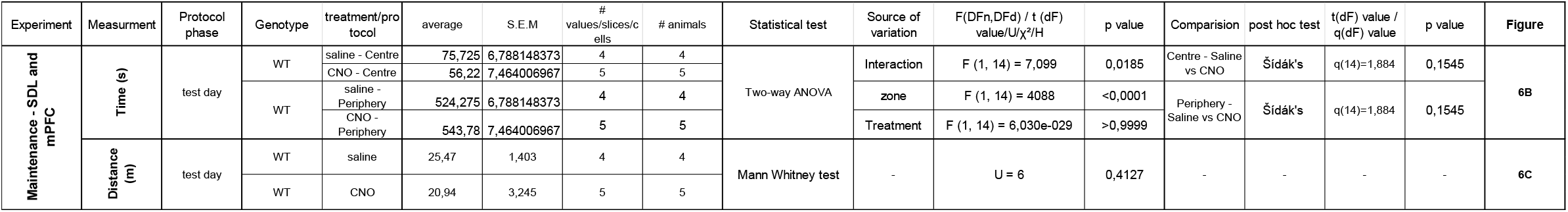
Statistics Table Figure 6.

## Notes

### Competing Interest Statement

The authors have declared no competing interest.

### Summary of Updates

We have done some corrections in the main manuscript

## REFERENCES

Akhlaghpour, H., Wiskerke, J., Choi, J.Y., Taliaferro, J.P., Au, J., Witten, I.B., 2016. Dissociated sequential activity and stimulus encoding in the dorsomedial striatum during spatial working memory. eLife 5, e19507. 10.7554/eLife.19507

Armbruster, B.N., Li, X., Pausch, M.H., Herlitze, S., Roth, B.L., 2007. Evolving the lock to fit the key to create a family of G protein-coupled receptors potently activated by an inert ligand. Proc. Natl. Acad. Sci. 104, 5163–5168. 10.1073/pnas.0700293104

Baddeley, A., 2000. The episodic buffer: a new component of working memory? Trends Cogn. Sci. 4, 417–423. 10.1016/S1364-6613(00)01538-2

Baddeley, A., 1986. Working memory., Working memory. Clarendon Press/Oxford University Press, New York, NY, US.

Baddeley, A.D., Hitch, G., 1974. Working Memory, in: Psychology of Learning and Motivation. Elsevier, pp. 47–89. 10.1016/S0079-7421(08)60452-1

Balleine, B.W., Dickinson, A., 1998. Goal-directed instrumental action: contingency and incentive learning and their cortical substrates. Neuropharmacology 37, 407–419. 10.1016/S0028-3908(98)00033-1

Balleine, B.W., O’Doherty, J.P., 2010. Human and Rodent Homologies in Action Control: Corticostriatal Determinants of Goal-Directed and Habitual Action. Neuropsychopharmacology 35, 48–69. 10.1038/npp.2009.131

Bouton, M.E., 2021. Context, attention, and the switch between habit and goal-direction in behavior. Learn. Behav. 49, 349–362. 10.3758/s13420-021-00488-z

Coutureau, E., Parkes, S.L., 2018. Cortical Determinants of Goal-Directed Behavior, in: Goal-Directed Decision Making. Elsevier, pp. 179–197. 10.1016/B978-0-12-812098-9.00008-5

Delatour, B., Gisquet-Verrier, P., 2000. Functional role of rat prelimbic-infralimbic cortices in spatial memory: evidence for their involvement in attention and behavioural flexibility. Behav. Brain Res. 109, 113–128. 10.1016/S0166-4328(99)00168-0

Dickinson, A., 1994. Instrumental Conditioning, in: Animal Learning and Cognition. Elsevier, pp. 45–79. 10.1016/B978-0-08-057169-0.50009-7

Dickinson, A., Balleine, B., Watt, A., Gonzalez, F., Boakes, R.A., 1995. Motivational control after extended instrumental training. Anim. Learn. Behav. 23, 197–206. 10.3758/BF03199935

Eichenbaum, H., 2002. The Cognitive Neuroscience of Memory: An Introduction, 1st ed. Oxford University Press New York. 10.1093/acprof:oso/9780195141740.001.0001

Euston, D.R., Gruber, A.J., McNaughton, B.L., 2012. The Role of Medial Prefrontal Cortex in Memory and Decision Making. Neuron 76, 1057–1070. 10.1016/j.neuron.2012.12.002

Funahashi, S., 2017. Working Memory in the Prefrontal Cortex. Brain Sci. 7, 49. 10.3390/brainsci7050049

Gisquet-Verrier, P., Delatour, B., 2006. The role of the rat prelimbic/infralimbic cortex in working memory: Not involved in the short-term maintenance but in monitoring and processing functions. Neuroscience 141, 585–596. 10.1016/j.neuroscience.2006.04.009

Graybiel, A.M., 1998. The basal ganglia and chunking of action repertoires. Neurobiol. Learn. Mem. 70, 119–136. 10.1006/nlme.1998.3843

Hasz, B.M., Redish, A.D., 2018. Deliberation and Procedural Automation on a Two-Step Task for Rats. Front. Integr. Neurosci. 12, 30. 10.3389/fnint.2018.00030

Hilario, M., Holloway, T., Jin, X., Costa, R.M., 2012a. Different dorsal striatum circuits mediate action discrimination and action generalization. Eur. J. Neurosci. 35, 1105–1114. 10.1111/j.1460-9568.2012.08073.x

Hilario, M., Holloway, T., Jin, X., Costa, R.M., 2012b. Different dorsal striatum circuits mediate action discrimination and action generalization. Eur. J. Neurosci. 35, 1105–1114. 10.1111/j.1460-9568.2012.08073.x

James, A., Reynaud-Bouret, P., Mezzadri, G., Sargolini, F., Bethus, I., Muzy, A., 2023. Strategy inference during learning via cognitive activity-based credit assignment models. Sci. Rep. 13, 9408. 10.1038/s41598-023-33604-2

Killcross, S., 2003. Coordination of Actions and Habits in the Medial Prefrontal Cortex of Rats. Cereb. Cortex 13, 400–408. 10.1093/cercor/13.4.400

Logie, R., Camos, V., Cowan, N. (Eds.), 2020. Working Memory: The state of the science, 1st ed. Oxford University Press. 10.1093/oso/9780198842286.001.0001

Miller, E.K., 2013. The “working” of working memory. Dialogues Clin. Neurosci. 15, 411–418. 10.31887/DCNS.2013.15.4/emiller

Monsell, S., Graham, B., 2021. Role of verbal working memory in rapid procedural acquisition of a choice response task. Cognition 214, 104731. 10.1016/j.cognition.2021.104731

Oberauer, K., 2006. Is the focus of attention in working memory expanded through practice? J. Exp. Psychol. Learn. Mem. Cogn. 32, 197–214. 10.1037/0278-7393.32.2.197

Packard, M.G., Knowlton, B.J., 2002. Learning and memory functions of the Basal Ganglia. Annu. Rev. Neurosci. 25, 563–593. 10.1146/annurev.neuro.25.112701.142937

Ragozzino, M.E., Detrick, S., Kesner, R.P., 1999. Involvement of the Prelimbic–Infralimbic Areas of the Rodent Prefrontal Cortex in Behavioral Flexibility for Place and Response Learning. J. Neurosci. 19, 4585–4594. 10.1523/JNEUROSCI.19-11-04585.1999

Sakai, K., 2008. Task Set and Prefrontal Cortex. Annu. Rev. Neurosci. 31, 219–245. 10.1146/annurev.neuro.31.060407.125642

Smith, K.S., Graybiel, A.M., 2013. A Dual Operator View of Habitual Behavior Reflecting Cortical and Striatal Dynamics. Neuron 79, 361–374. 10.1016/j.neuron.2013.05.038

Spellman, T., Rigotti, M., Ahmari, S.E., Fusi, S., Gogos, J.A., Gordon, J.A., 2015. Hippocampal–prefrontal input supports spatial encoding in working memory. Nature 522, 309–314. 10.1038/nature14445

Tonegawa, S., Morrissey, M.D., Kitamura, T., 2018. The role of engram cells in the systems consolidation of memory. Nat. Rev. Neurosci. 19, 485–498. 10.1038/s41583-018-0031-2

Tunes, G.C., Fermino De Oliveira, E., Vieira, E.U., Caetano, M.S., Cravo, A.M., Bussotti Reyes, M., 2022. Time encoding migrates from prefrontal cortex to dorsal striatum during learning of a self-timed response duration task. eLife 11, e65495. 10.7554/eLife.65495

Uylings, H.B.M., Groenewegen, H.J., Kolb, B., 2003. Do rats have a prefrontal cortex? Behav. Brain Res. 146, 3–17. 10.1016/j.bbr.2003.09.028

Vandierendonck, A., 2016. A Working Memory System With Distributed Executive Control. Perspect. Psychol. Sci. 11, 74–100. 10.1177/1745691615596790

Vogel, P., Hahn, J., Duvarci, S., Sigurdsson, T., 2022. Prefrontal pyramidal neurons are critical for all phases of working memory. Cell Rep. 39, 110659. 10.1016/j.celrep.2022.110659

Wilhelm, M., Sych, Y., Fomins, A., Alatorre Warren, J.L., Lewis, C., Serratosa Capdevila, L., Boehringer, R., Amadei, E.A., Grewe, B., O’Connor, E.C., Hall, B.J., Helmchen, F., 2023. Striatum-projecting prefrontal cortex neurons support working memory maintenance. Nat. Commun. 14, 7016. 10.1038/s41467-023-42777-3

Yang, S.-T., Shi, Y., Wang, Q., Peng, J.-Y., Li, B.-M., 2014. Neuronal representation of working memory in the medial prefrontal cortex of rats. Mol. Brain 7, 61. 10.1186/s13041-014-0061-2

Yin, H.H., Knowlton, B.J., Balleine, B.W., 2004. Lesions of dorsolateral striatum preserve outcome expectancy but disrupt habit formation in instrumental learning. Eur. J. Neurosci. 19, 181–189. 10.1111/j.1460-9568.2004.03095.x

Yin, H.H., Ostlund, S.B., Knowlton, B.J., Balleine, B.W., 2005. The role of the dorsomedial striatum in instrumental conditioning. Eur. J. Neurosci. 22, 513–523. 10.1111/j.1460-9568.2005.04218.x

Yoon, T., Okada, J., Jung, M.W., Kim, J.J., 2008. Prefrontal cortex and hippocampus subserve different components of working memory in rats. Learn. Mem. 15, 97–105. 10.1101/lm.850808

