## supplementary figures and tables for "Cortico-striatal dynamics across working memory stages"

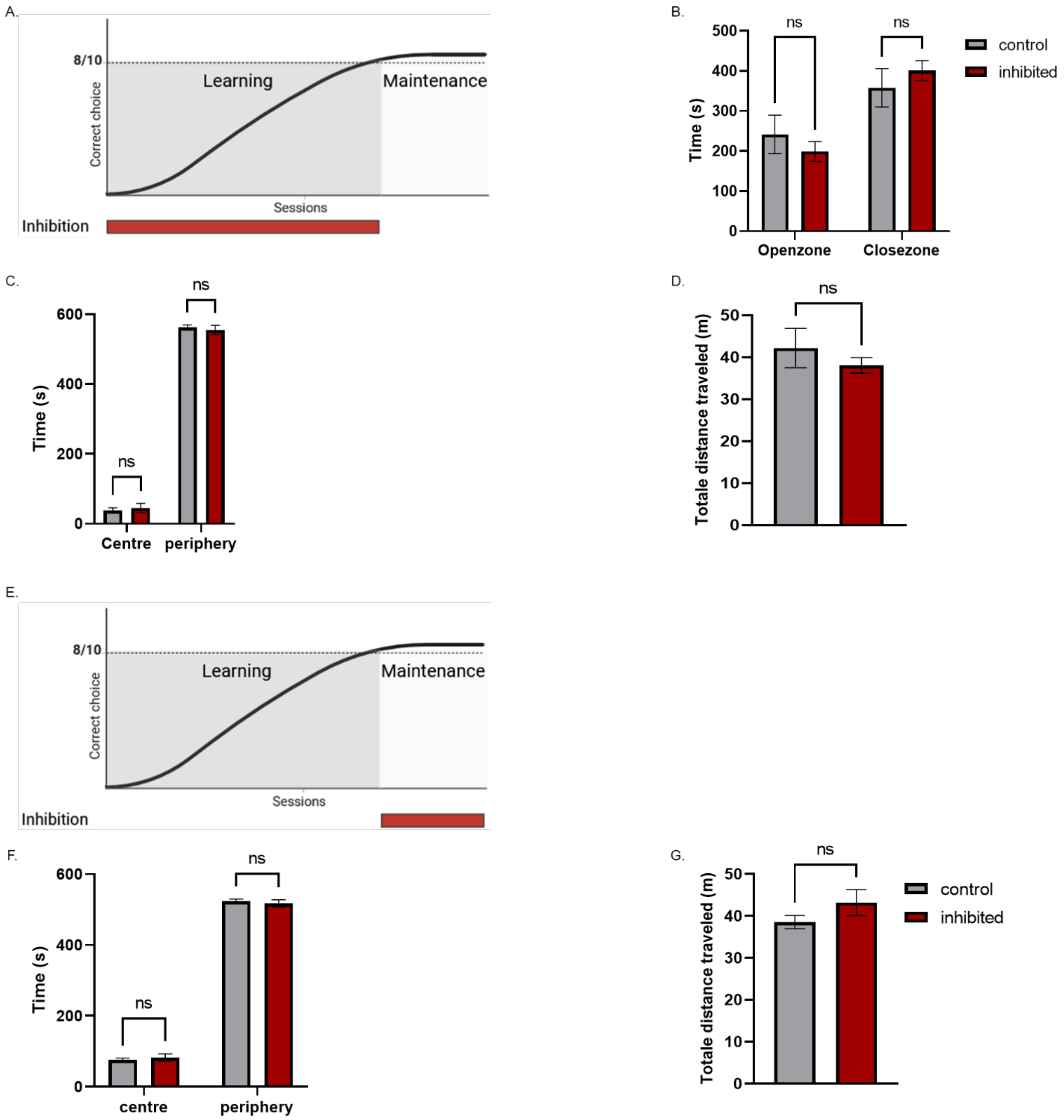


**Figure S1: Evaluation of locomotion and anxiety in mice with inhibition of the DLS.**
**A**. Experimental design of inhibition during the learning phase of DNMP task. **B**. Average time spent by animals in the open and closed areas of the O-maze during ten minutes. **C**. Average time spent by animals in the central and peripheral zones of the open field during ten minutes. **D**. Total distance covered by animals in an open field. **E**. Experimental design of inhibition during the maintenance phase of DNMP task. **F**. Average time spent by animals in the central and peripheral zones of the open field during ten minutes. **G**. Total distance covered by animals in an open field. (control n= 5, inhibited n= 6; B: Two-way anova, open zone: control x inhibited, “ns” p=0,6535, close zone: control x inhibited, “ns” p= 0,6535; C: Two-way anova, centre: control x inhibited, “ns” p= 0,8873, periphery: control x inhibited, “ns” p= 0,8873; D: Unpaired t test, control x inhibited, “ns” p= 0,4028; control n= 7, inhibited n= 7; F: Two-way anova, centre: control x inhibited, “ns” p= 0,8007, periphery: control x inhibited, “ns” p= 0,8007; G : Paired t test, control x inhibited, “ns” p= 0,2463).

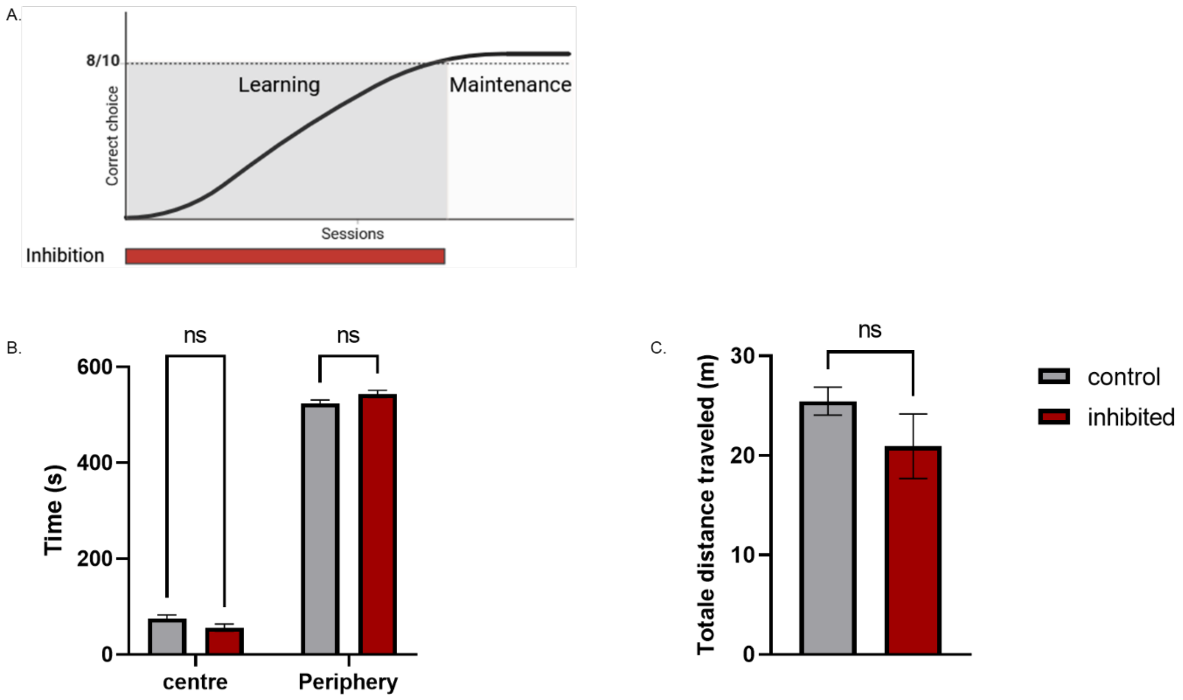


**Figure S2: Evaluation of locomotion and anxiety in mice with inhibition of mPFC and the DLS.**

**A**. Experimental design of inhibition during the maintenance phase of DNMP task. **B.** Average time spent by animals in the central and peripheral zones of the open field during ten minutes. **C**. Total distance covered by animals in an open field. (control n= 5, inhibited n= 4; B: Two-way anova, centre: control x inhibited, “ns” p= 0,1545, periphery: control x inhibited, “ns” p= 0,1545; C: Mann-Whitney test, control x inhibited, “ns” p= 0,4127).

### Statistics Table Figure 1

| Experiment | Measurment | Protocol phase | Genotype | treatment/protocol | average | S.E.M | # values/slices/cells | # animals | Statistical test | Source of variation | F(DFn,DFd) / t (dF) value/U/$\chi^2$/H | p value | Comparision | post hoc test | t(dF) value / q(dF) value | p value | Figure |
|---|---|---|---|---|---|---|---|---|---|---|---|---|---|---|---|---|---|
| Devaluation - Learning | criterion | test day | WT | devalued | 0,5125 | 0,05489 | 8 | 8 | one sample t-test | devalued vs criterion | t=3,416, df=7 | 0,0112 | - | - | - | - | 1D |
| | | | criterion | | 0,7 | | | | | | | | | | | | |
| | | | WT | non-devalued | 0,75 | 0,04226 | 8 | 8 | | non-devalued vs criterion | t=1,183, df=7 | 0,2753 | - | - | - | - | |
| | | | criterion | | 0,7 | | | | | | | | | | | | |
| | mean of test day | test day | WT | TTC | 0,775 | 0,03134 | 8 | 8 | One-way ANOVA | Treatment | F (1,717, 12,02) = 12,54 | 0,0015 | TTC vs devalued | Holm-Šídák's | q(7)=5,475 | 0,0028 | 1E |
| | | | WT | devalued | 0,5125 | 0,05489 | 8 | 8 | | | | | TTC vs non-devalued | | q(7)=0,4472 | 0,6682 | |
| | | | WT | non-devalued | 0,75 | 0,04226 | 8 | 8 | | | | | devalued vs non-devalued | | q(7)=3,494 | 0,02 | |
| Devaluation - Maintenance | criterion | test day | WT | devalued | 0,8 | 0,04924 | 12 | 12 | one sample Wilcoxon test | devalued vs criterion | - | >0,9999 | - | - | - | - | 1G |
| | | | criterion | | 0,8 | | | | | | | | | | | | |
| | | | WT | non-devalued | 0,8 | 0,04767 | 12 | 12 | | non-devalued vs criterion | - | >0,9999 | - | - | - | - | |
| | | | criterion | | 0,8 | | | | | | | | | | | | |
| | mean of test day | test day | WT | TTC | 0,8111 | 0,0226 | 12 | 12 | Friedman test | - | - | 0,7008 | TTC vs devalued | Dunn's | - | >0,9999 | 1H |
| | | | WT | devalued | 0,8 | 0,04924 | 12 | 12 | | | | | TTC vs non-devalued | | - | >0,9999 | |
| | | | WT | non-devalued | 0,8 | 0,04767 | 12 | 12 | | | | | devalued vs non-devalued | | - | >0,9999 | |

### Statistics Table Figure 2

| Experiment | Measurment | Protocol phase | Genotype | treatment/protocol | average | S.E.M | # values/slices/cells | # animals | Statistical test | Source of variation | F(DFn,DFd) / t (dF) value/U/$\chi^2$/H | p value | Comparision | post hoc test | t(dF) value / q(dF) value | p value | Figure |
|---|---|---|---|---|---|---|---|---|---|---|---|---|---|---|---|---|---|
| Learning - mPFC | criterion | S9 S10 S11 | WT | saline | 0,8 | 0,02854 | 10 | 10 | one sample t-test | saline vs criterion | t=0,000, df=9 | >0,9999 | - | - | - | - | 2C |
| | | | criterion | | 0,8 | | | | | | | | | | | | |
| | | | WT | CNO | 0,6233 | 0,03786 | 10 | 10 | | CNO vs criterion | t=4,666, df=9 | 0,0012 | - | - | - | - | |
| | | | criterion | | 0,8 | | | | | | | | | | | | |
| | mean of test day | S9 S10 S12 | WT | saline | 0,8 | 0,02854 | 10 | 10 | Unpaired t test | saline vs CNO | t=3,726, df=18 | 0,0008 | - | - | - | - | 2D |
| | | | WT | CNO | 0,6233 | 0,03786 | 10 | 10 | | | | | | | | | |
| Maintenance - mPFC | criterion | test day | WT | saline | 0,775 | 0,03708 | 16 | 16 | one sample t-test | saline vs criterion | t=0,6742, df=15 | 0,5104 | - | - | - | - | 2F |
| | | | criterion | | 0,8 | | | | | | | | | | | | |
| | | | WT | CNO | 0,7813 | 0,02772 | 16 | 16 | | CNO vs criterion | t=0,6765, df=15 | 0,509 | - | - | - | - | |
| | | | criterion | | 0,8 | | | | | | | | | | | | |
| | mean of test day | test day | WT | TTC | 0,7875 | 0,02295 | 16 | 16 | RM One-way ANOVA | treatment | F (1,465, 21,98) = 0,04931 | 0,9064 | TTC vs saline | Holm-Šídák's | q(15)=0,3661 | 0,9779 | 2G |
| | | | WT | saline | 0,775 | 0,03708 | 16 | 16 | | | | | TTC vs CNO | Holm-Šídák's | q(15)=0,1930 | 0,9779 | |
| | | | WT | CNO | 0,7813 | 0,02772 | 16 | 16 | | | | | saline vs CNO | Holm-Šídák's | q(15)=0,1240 | 0,9779 | |

### Statistics Table Figure 3

| Experiment | Measurment | Protocol phase | Genotype | treatment/protocol | average | S.E.M | # values/slices/cells | # animals | Statistical test | Source of variation | F(DFn,DFd) / t (dF) value/U/$\chi^2$/H | p value | Comparision | post hoc test | t(dF) value / q(dF) value | p value | Figure |
|---|---|---|---|---|---|---|---|---|---|---|---|---|---|---|---|---|---|
| Learning - SDL | criterion | T-3 T-2 T-1 | WT | saline | 0,8533 | 0,017 | 5 | 5 | one sample t-test | saline vs criterion | t=3,138, df=4 | 0,0349 | - | - | - | - | 3C |
| | | | criterion | | 0,8 | | | | | | | | | | | | |
| | | | WT | CNO | 0,8778 | 0,01648 | 6 | 6 | | CNO vs criterion | t=4,719, df=5 | 0,0052 | - | - | - | - | |
| | | | criterion | | 0,8 | | | | | | | | | | | | |
| | mean of test day | T-3 T-2 T-2 | WT | saline | 0,8533 | 0,017 | 5 | 5 | Unpaired t test | saline vs CNO | t=1,026, df=9 | 0,3316 | - | - | - | - | 3D |
| | | | WT | CNO | 0,8778 | 0,01648 | 6 | 6 | | | | | | | | | |
| | mean of test day | test day | WT | TTC | 0,8533 | 0,017 | 5 | 5 | RM One-way ANOVA | treatment | F (1,604, 6,414) = 1,155 | 0,3578 | TTC vs devalued | Holm-Šídák's | q(4)=1,548 | 0,5659 | 3E |
| | | | WT | saline / devalued | 0,78 | 0,07348 | 5 | 5 | | | | | TTC vs non-devalued | Holm-Šídák's | q(4)=2,429 | 0,3054 | |
| | | | WT | saline / non-devalued | 0,78 | 0,05831 | 5 | 5 | | | | | devalued vs non-devalued | Holm-Šídák's | q(4)=0,000 | >0,9999 | |
| | mean of test day | test day | WT | TTC | 0,8778 | 0,01648 | 6 | 6 | RM One-way ANOVA | treatment | F (1,512, 7,560) = 27,59 | 0,0005 | TTC vs devalued | Holm-Šídák's | q(5)=7,050 | 0,0027 | 3F |
| | | | WT | CNO / devalued | 0,5167 | 0,05426 | 6 | 6 | | | | | TTC vs non-devalued | Holm-Šídák's | q(5)=1,136 | 0,3075 | |
| | | | WT | CNO / non-devalued | 0,8333 | 0,04216 | 6 | 6 | | | | | devalued vs non-devalued | Holm-Šídák's | q(5)=4,842 | 0,0094 | |
| Maintenance - SDL | criterion | test day | WT | saline | 0,875 | 0,02787 | 12 | 12 | one sample t-test | saline vs criterion | t=2,691, df=11 | 0,021 | - | - | - | - | 3H |
| | | | criterion | | 0,8 | | | | | | | | | | | | |
| | | | WT | CNO | 0,7833 | 0,03445 | 12 | 12 | | CNO vs criterion | t=0,4838, df=11 | 0,638 | - | - | - | - | |
| | | | criterion | | 0,8 | | | | | | | | | | | | |
| | mean of test day | test day | WT | TTC | 0,8528 | 0,008663 | 12 | 12 | RM One-way ANOVA | treatment | F (1,713, 18,84) = 3,469 | 0,0583 | TTC vs saline | Holm-Šídák's | q(11)=0,7790 | 0,4524 | 3I |
| | | | WT | saline | 0,875 | 0,02787 | 12 | 12 | | | | | TTC vs CNO | Holm-Šídák's | q(11)=1,853 | 0,1736 | |
| | | | WT | CNO | 0,7833 | 0,03445 | 12 | 12 | | | | | Saline vs CNO | Holm-Šídák's | q(11)=2,200 | 0,1429 | |
| | mean of test day | test day | WT | TTC | 0,8528 | 0,008663 | 12 | 12 | RM One-way ANOVA | treatment | F (1,401, 15,41) = 29,62 | <0,0001 | TTC vs devalued - saline | Holm-Šídák's | q(11)=3,089 | 0,1182 | 3J |
| | | | WT | devalued - saline | 0,775 | 0,03718 | 12 | 12 | | | | | TTC vs devalued - CNO | Holm-Šídák's | q(11)=12,76 | <0,0001 | |
| | | | WT | devalued - CNO | 0,5333 | 0,03553 | 12 | 12 | | | | | devalued - Saline vs devalued - [illegible] | Holm-Šídák's | q(11)=6,139 | 0,0031 | |

### Statistics Table Figure 4

| Experiment | Measurment | Protocol phase | Genotype | treatment/protocol | average | S.E.M | # values/slices/cells | # animals | Statistical test | Source of variation | F(DFn,DFd) / t (dF) value/U/$\chi^2$/H | p value | Comparision | post hoc test | t(dF) value / q(dF) value | p value | Figure |
|---|---|---|---|---|---|---|---|---|---|---|---|---|---|---|---|---|---|
| Maintenance - SDL and mPFC | criterion | test day | WT | saline | 0,8571 | 0,03689 | 7 | 7 | one sample t-test | saline vs criterion | t=1,549, df=6 | 0,1723 | - | - | - | - | 4C |
| | | | criterion | | 0,8 | | | | | | | | | | | | |
| | | | WT | CNO | 0,6429 | 0,03689 | 7 | 7 | | CNO vs criterion | t=4,260, df=6 | 0,0053 | - | - | - | - | |
| | | | criterion | | 0,8 | | | | | | | | | | | | |
| | mean of test day | test day | WT | TTC | 0,8476 | 0,0123 | 7 | 7 | RM One-way ANOVA | treatment | F (1,914, 11,48) = 16,62 | 0,0004 | TTC vs saline | Holm-Šídák's | q(6)=0,2487 | 0,8119 | 4D |
| | | | WT | saline | 0,8571 | 0,03689 | 7 | 7 | | | | | TTC vs CNO | Holm-Šídák's | q(6)=4,943 | 0,0078 | |
| | | | WT | CNO | 0,6429 | 0,03689 | 7 | 7 | | | | | Saline vs CNO | Holm-Šídák's | q(6)=4,666 | 0,0078 | |

### Statistics Table Figure 5

| Experiment | Measurment | Protocol phase | Genotype | treatment/protocol | average | S.E.M | # values/slices/cells | # animals | Statistical test | Source of variation | F(DFn,DFd) / t (dF) value/U/$\chi^2$/H | p value | Comparision | post hoc test | t(dF) value / q(dF) value | p value | Figure |
|---|---|---|---|---|---|---|---|---|---|---|---|---|---|---|---|---|---|
| SDL - Learning | Time (s) | test day | WT | saline - Openzone | 242,04 | 47,88077485 | 5 | 5 | Two-way ANOVA | Interaction | F (1, 18) = 1,415 | 0,2497 | Openzone - Saline vs CNO | Šídák's | q(18)=0,8410 | 0,6535 | 5B |
| | | | | CNO - Openzone | 198,9666667 | 24,85505538 | 6 | 6 | | | | | | | | | |
| | | | WT | saline - Closezone | 357,96 | 47,88077485 | 5 | 5 | | zone | F (1, 18) = 19,27 | 0,0004 | Closezone - Saline vs CNO | Šídák's | q(18)=0,8410 | 0,6535 | |
| | | | | CNO - Closezone | 401,0333333 | 24,85505538 | 6 | 6 | | Treatment | F (1, 18) = 6,159e-031 | >0,9999 | | | | | |
| | Time (s) | test day | WT | saline - Centre | 37,96 | 7,613645644 | 5 | 5 | Two-way ANOVA | Interaction | F (1, 18) = 0,3893 | 0,5405 | Centre - Saline vs CNO | Šídák's | q(18)=0,4412 | 0,8873 | 5C |
| | | | | CNO - Centre | 45,06666667 | 13,17737118 | 6 | 6 | | | | | | | | | |
| | | | WT | saline - Periphery | 562,04 | 7,613645644 | 5 | 5 | | zone | F (1, 18) = 2060 | <0,0001 | Periphery - Saline vs CNO | Šídák's | q(18)=0,4412 | 0,8873 | |
| | | | | CNO - Periphery | 554,9333333 | 13,17737118 | 6 | 6 | | Treatment | F (1, 18) = 0,000 | >0,9999 | | | | | |
| | Distance (m) | test day | WT | saline | 42,21 | 4,681 | 5 | 5 | Unpaired t test | saline vs CNO | t=0,8779, df=9 | 0,4028 | - | - | - | - | 5D |
| | | | WT | CNO | 38,08 | 1,849 | 6 | 6 | | | | | | | | | |
| SDL - Maintenance | Time (s) | test day | WT | saline - Centre | 75,55714286 | 5,160696573 | 7 | 7 | Two-way ANOVA | Interaction | F (1, 24) = 0,7221 | 0,4039 | Centre - Saline vs CNO | Šídák's | q(24)=0,6009 | 0,8007 | 5F |
| | | | | CNO - Centre | 82,45714286 | 10,25837305 | 7 | 7 | | | | | | | | | |
| | | | WT | saline - Periphery | 524,4428571 | 5,160696573 | 7 | 7 | | zone | F (1, 24) = 2963 | <0,0001 | Periphery - Saline vs CNO | Šídák's | q(24)=0,6009 | 0,8007 | |
| | | | | CNO - Periphery | 517,5428571 | 10,25837305 | 7 | 7 | | Treatment | F (1, 24) = 0,000 | >0,9999 | | | | | |
| | Distance (m) | test day | WT | saline | 38,54 | 1,605 | 7 | 7 | Paired t test | saline vs CNO | t=1,285, df=6 | 0,2463 | - | - | - | - | 5G |
| | | | WT | CNO | 43,21 | 3,116 | 7 | 7 | | | | | | | | | |

### Statistics Table Figure 6

| Experiment | Measurment | Protocol phase | Genotype | treatment/protocol | average | S.E.M | # values/slices/cells | # animals | Statistical test | Source of variation | F(DFn,DFd) / t (dF) value/U/$\chi^2$/H | p value | Comparision | post hoc test | t(dF) value / q(dF) value | p value | Figure |
|---|---|---|---|---|---|---|---|---|---|---|---|---|---|---|---|---|---|
| Maintenance - SDL and mPFC | Time (s) | test day | WT | saline - Centre | 75,725 | 6,788148373 | 4 | 4 | Two-way ANOVA | Interaction | F (1, 14) = 7,099 | 0,0185 | Centre - Saline vs CNO | Šídák's | q(14)=1,884 | 0,1545 | 6B |
| | | | | CNO - Centre | 56,22 | 7,464006967 | 5 | 5 | | | | | | | | | |
| | | | WT | saline - Periphery | 524,275 | 6,788148373 | 4 | 4 | | zone | F (1, 14) = 4088 | <0,0001 | Periphery - Saline vs CNO | Šídák's | q(14)=1,884 | 0,1545 | |
| | | | | CNO - Periphery | 543,78 | 7,464006967 | 5 | 5 | | Treatment | F (1, 14) = 6,030e-029 | >0,9999 | | | | | |
| | Distance (m) | test day | WT | saline | 25,47 | 1,403 | 4 | 4 | Mann Whitney test | - | U = 6 | 0,4127 | - | - | - | - | 6C |
| | | | WT | CNO | 20,94 | 3,245 | 5 | 5 | | | | | | | | | |
